# Multiple triggers converge to preferential effector coupling in the CB_2_R through a complex allosteric communication network

**DOI:** 10.1101/2024.02.27.582279

**Authors:** Adrian Morales-Pastor, Tamara Miljuš, Tomasz Stępniewski, Vicente Ledesma, Franziska M. Heydenreich, Tilman Flock, Bianca Plouffe, Christian Le Gouill, Jean Duchaine, Wolfgang Guba, Arne C. Rufer, Uwe Grether, Michel Bouvier, Dmitry B. Veprintsev, Jana Selent

## Abstract

G protein-coupled receptors are important therapeutic drug targets for a wide range of diseases. Their ability to preferentially engage specific signaling pathways over others can be exploited to design drugs that target only disease-associated pathways leading to an improved safety profile. However, the underlying molecular mechanisms for preferential pathway engagement are complex and remain largely elusive. To elucidate the multifaceted actions at the receptor level that lead to preferential coupling, we employ a combination of techniques. Our approach integrates systematic mutagenesis of the CB_2_R and comprehensive profiling of Gαi2 and β-arrestin1 engagements with computer simulations to track mutant-induced impacts on receptor dynamics. Most importantly, our research discloses multiple triggers on a complex allosteric communication network (ACN) that converge to preferential CB_2_R coupling by modulating evolutionary conserved motifs (e.g., CWxP, NPxxY, sodium binding site). Potent triggers for a preferential Gαi2 response exhibit high levels of connectivity and are located in proximity to connections with high information transmission. Our insights highlight the complexity of GPCR signaling and can guide the rational design of drug candidates tailored to evoke specific functional responses that can enhance the precision and efficacy of therapeutic interventions.

## Introduction

G protein-coupled receptors (GPCRs) are the largest family of cell-surface receptors in humans^1^. They are involved in virtually every physiological process which makes them invaluable targets for drug development. The cannabinoid receptor 2 (CB_2_R) is a GPCR that plays a crucial role in numerous physiological processes including immune response and inflammation^2^. This makes the CB_2_R a promising therapeutic target for a variety of diseases, such as chronic pain, neuroinflammation, and cancer^3,4^. Currently, the only FDA approved drugs acting on CB_2_R are phytocannabinoids including Δ9-THC or its synthetic analogues. However, these compounds are non-selective agonist of cannabinoid receptors and cause psychoactive effects via CB1R^4^. Additionally, they also cause adverse cardio-vascular liability in a dose-dependent matter^5^ via CB1R. In contrast, CB_2_R agonists seem to be devoid of adverse side effects. There are over 20 selective CB_2_R agonists in clinical trials for a variety of inflammation conditions that show good safety profiles. Unfortunately, these compounds have not yet transformed into approved therapeutics, reflecting their elusive therapeutic effect. A potential reason for this is a broad distribution of CB_2_R in different tissues, as well “on demand” engagement of the endocannabinoid system^6^, making the development of systemically applied drugs challenging. More selective drugs are needed to control the endocannabinoid system with greater precision. A promising approach to improve the specificity of drug action as well as modulate its efficacy and to reduce potential side effects, is to exploit the concept of signaling bias. Hereby, a drug preferentially activates specific downstream signaling pathways over ones that are related to unwanted side effects^7–9^.

Recent work provides evidence for the relevance of allosteric communication and implicated networks in GPCR function and signaling bias^10,11^. Allosteric communication networks (ACNs) serve as dynamic infrastructure that facilitates communication between remote regions within GPCRs, connecting critical sites like the ligand-binding site to the intracellular effector coupling site. However, the details of how ACNs drive preferential engagement of a specific effector protein remain largely elusive. A main challenge of studying allostery in GPCRs is to monitor ACNs and their alterations related to specific signaling conditions at appropriate spatio-temporal resolution. Molecular dynamics (MD) simulations have emerged as an invaluable tool for complementing static structural, biophysical and biochemical data^12–14^ to access the proper resolution levels. They allow for the exploration of protein dynamics at an atomic level, offering insights into the conformational changes at microsecond timescale and corresponding intramolecular networks under conditions of signaling bias^15,16^. Knowledge of the amino acid networks involved in this allosteric communication can not only advance our understanding of GPCR signaling bias but also boost the design of biased ligands with a more precise therapeutic profile.

In this study, we dissect the allosteric communication networks in the CB_2_R, combining high-throughput state-of-the-art computational and cell-based assay methods. In a systematic study, we first probe the perturbation of each individual residue within the primary structure of the CB_2_ through mutagenesis combined with G protein activation and β-arrestin1 (βarr1) recruitment. This information is integrated with time-resolved molecular dynamics of the mutation-induced alteration of the CB_2_R ACNs using a machine learning pipeline. Most importantly, our study highlights multiple molecular mechanisms and how they interact with the main communication channels between the orthosteric ligand binding site and the intracellular binding effector interfaces via highly conserved residues. Of interest, mutations that preferentially trigger G protein coupling show high connectivity and are typically located closer to ligand-stabilized communication channels of high information transmission. These insights have important implications for the rational development of novel biased ligands targeting not only the orthosteric site but also allosteric sites in the CB_2_R with a therapeutic potential to treat inflammation and pain.

## Results

### Mutant position and sequence conservation impacts receptor cell-surface expression

First, we investigated 360 point mutations in the receptor to alanine or valine if the native amino acid was an alanine and its impact on cell-surface expression (see **Supplementary Data 1, Figure 1A-C**). We find that approximately 60% of all mutants exhibit a cell-surface expression within the wild-type (WT)-like range (i.e., 80% to 120% of WT expression) (**Figure 1B**). This finding suggests that most residue positions in CB_2_R display remarkable resilience in maintaining their cell-surface expression upon mutation to alanine/valine. Interestingly, approximately 30% of the mutants show a decrease in expression below the WT-like level, while only 10% of the mutants display an increase (**Figure 1B**). This observation indicates that expression-impacting mutants in the CB_2_R predominantly reduce cell-surface expression rather than increasing it. Such an effect can be attributed to mutation-induced misfolding and subsequent sequestration in the endoplasmic reticulum or the Golgi^17^.

**Figure 1.**
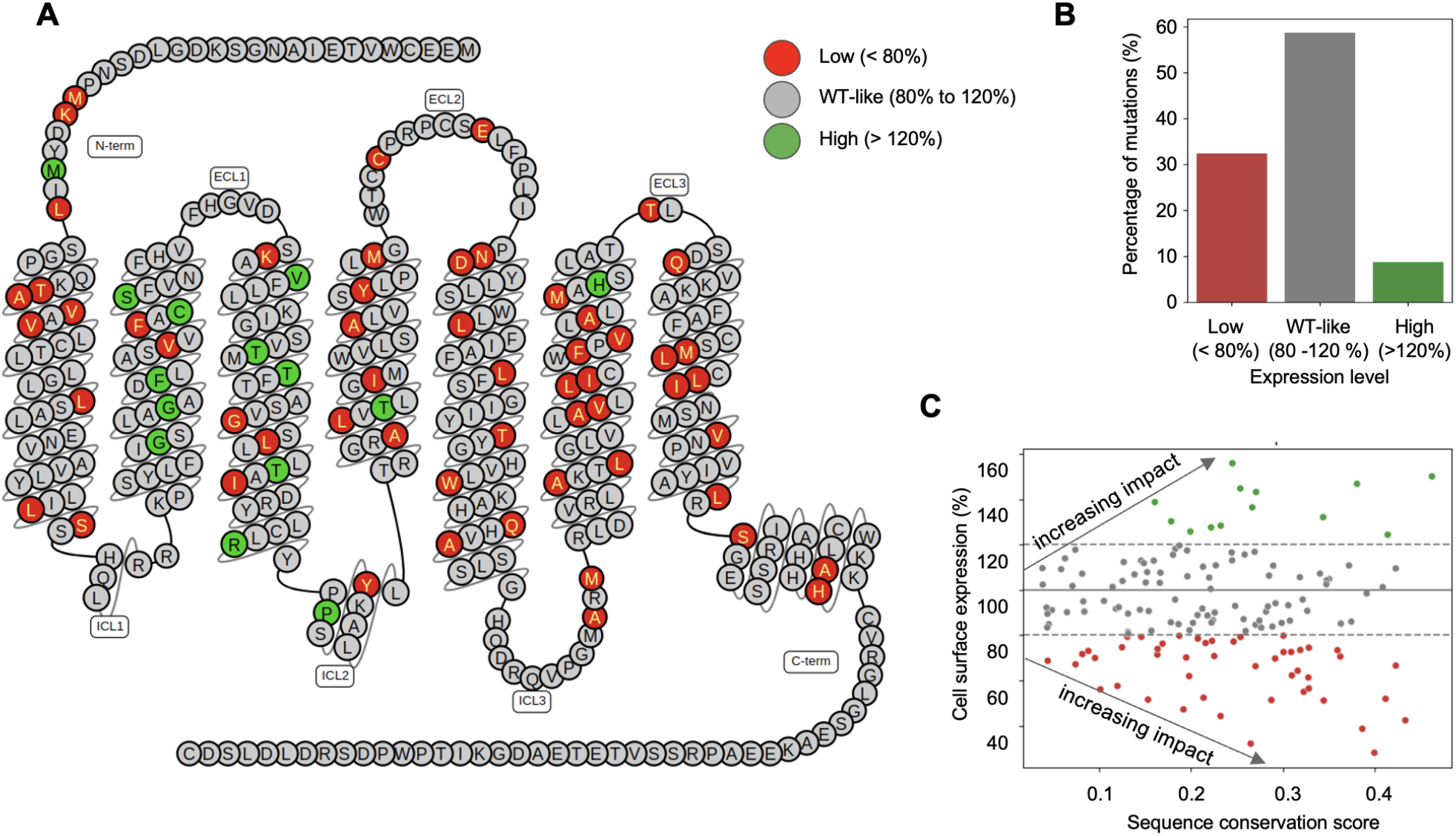
Surface expression and conservation. (**A**) CB_2_R snake plot showing the distribution of mutants with low, WT-like, and high surface expression. These categories correspond to less than 80% (red), between 80% and 120% (gray), and more than 120% (green) of the wild-type expression level. (**B**) Barplot showing the distribution of the three expression categories: low, WT-like and high. (**C**) Relationship between evolutionary conservation and the impact of mutations on cell-surface expression. The conservation score (Jensen-Shannon divergence) was computed for all residues in class A GPCRs. Residues with low conservation scores have the least effect on cell-surface expression, whereas high conservation scores can significantly impact cell-surface expression. Gray arrows indicate the increasing impact. WT surface expression level (100%) is shown with a solid horizontal gray line, while the two gray dashed lines delimit the boundaries of the WT-like expression level.

Of note, the distribution of expression groups across the receptor sequence reveals an interesting pattern (**Figure 1A**). Mutations that result in high expression levels in the CB_2_R tend to be preferentially located in transmembrane domains (TMs) 2 and 3 (green circles). On the other hand, mutants associated with low expression (red circles) are more evenly dispersed throughout the transmembrane core of the protein, with fewer occurrences in TM2 and TM3. This spatial association between expression groups suggests a potential functional significance of these specific transmembrane domains in modulating cell-surface expression levels in CB_2_R.

Furthermore, we studied the relationship between sequence conservation and cell-surface expression. The sequence conservation in the CB_2_R was computed based on the Jensen-Shannon Divergence (JSD) scoring method^18^ from a class A human GPCR alignment obtained from GPCRdb^19^. We observe that residues with lower conservation scores (< 0.2) show less impact on cell-surface expression with most mutants belonging to the WT-like group (grey points, **Figure 1C**). In contrast, residues with conservation scores > 0.2 yield larger expression deviations compared to the WT (arrows). This finding supports the widely accepted viewpoint that highly conserved residues are crucial for vital receptor functions, such as protein folding or trafficking to the cell membrane. Of interest are also residues that despite being highly conserved (conservation score > 0.4) show a WT-like expression level upon mutation (grey points). Those residues are likely involved in relevant receptor functions other than cell-surface expression,e.g. ligand binding, coupling to transducer proteins or signal transduction.

### Low mutational resilience of βarr1 recruitment results in preferential Gαi2 coupling

We also systematically assessed the effect of 360 point mutations within the CB_2_R on both Gαi2 coupling and βarr1 recruitment after stimulation with the agonist HU-210 using the ebBRET-based Effector Membrane Translocation Assay (EMTA) biosensor platform (**Supplementary Data 1**). Briefly, receptor-mediated Gαi2 activation was monitored through the translocation of a downstream effector Rap1GAP subunit to the plasma membrane, where it selectively interacts with activated Gi/o protein subfamily^20^. The same plasma membrane translocation principle is used to measure βarr1 recruitment to the receptor^17^.

For this large-scale analysis, we decided to focus on cases with a marked impact on coupling, i.e. the loss of Gαi2 or βarr1 engagement or both to obtain meaningful results. We defined four classes of receptor mutants (**Figure 2A**, see methods): (i) preferential Gαi2 coupling (PrefCoup_Gαi2_): no measurable βarr1 recruitment but preserved Gαi2 coupling (Emax > 50%), (ii) preferential βarr1 recruitment (PrefCoup_βarr1_): no measurable Gαi2 coupling but preserved βarr1 recruitment (Emax > 50%), (iii) loss of receptor function (NoCoup_Gαi2_βarr1_): no measurable signal for Gαi2 nor βarr1 and finally (iv) preserved coupling (Coup_Gαi2_βarr1_): mutations that preserve Gαi2 coupling and βarr1 recruitment with an Emax > 50%. Using this classification, we observe that 70% of mutants preserve coupling properties (Emax > 50%) to Gαi2 and βarr1, whereas 30% result in a loss of Gαi2, βarr1 or both (**Figure 2B**). Interestingly, our data indicate that coupling-impaired mutants predominantly induce the loss of βarr1 recruitment (19%), leading to preferential Gαi2 coupling, whereas Gαi2 coupling seems to be more robust to mutational alterations (only 2% specifically abrogated Gαi2 activation).

**Figure 2.**
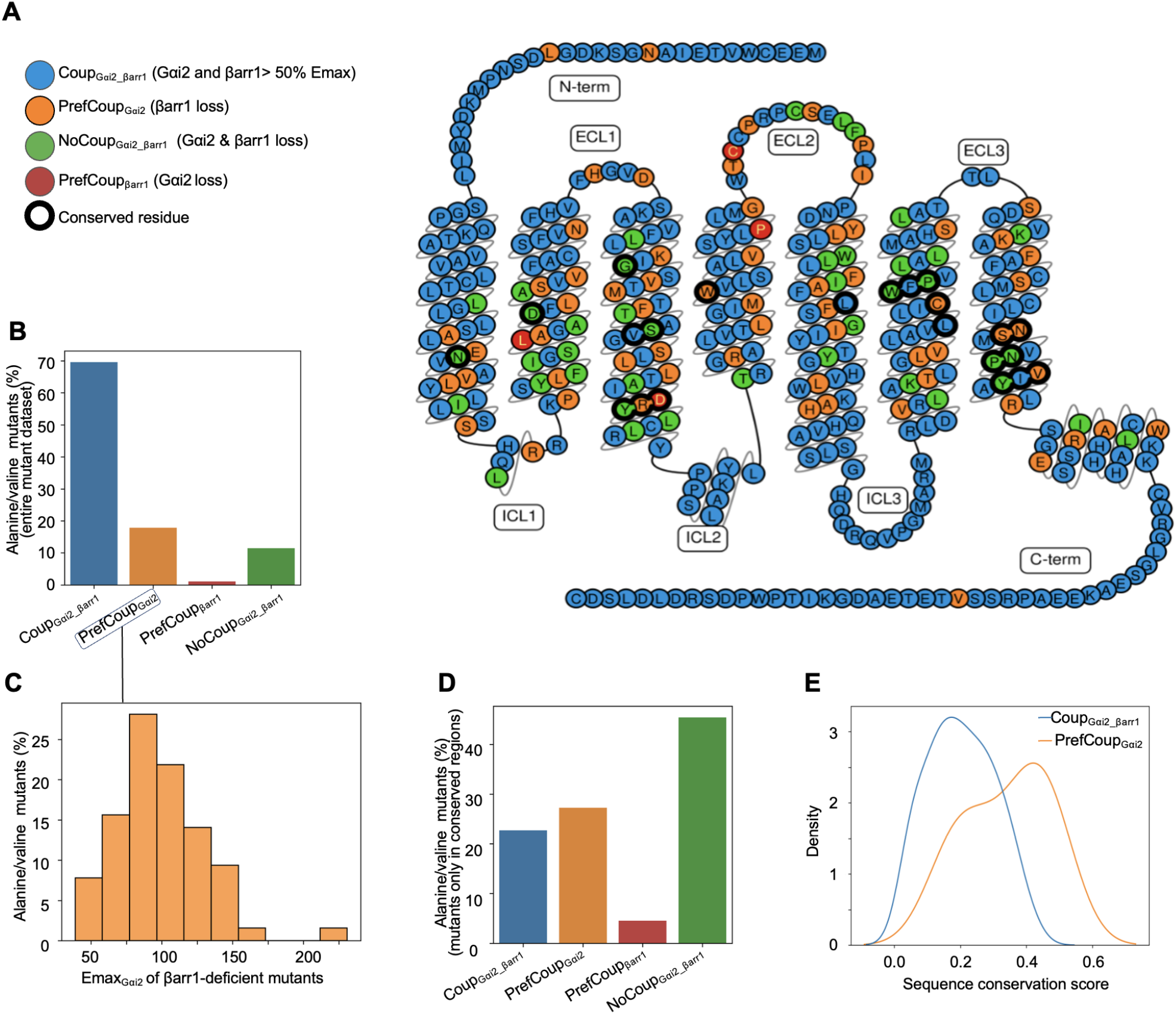
Coupling profiles and conservation. (**A**) Location of mutations in the CB_2_R classified as: Coup_Gαi2_βarr1_: preserved Gαi2 and βarr1 coupling (blue), PrefCoup_Gαi_: preferential Gαi2 coupling through a loss of βarr1 recruitment (orange), PrefCoup_βarr1_: preferential βarr1 recruitment through loss of Gαi2 coupling (red), and NoCoup_Gαi2_βarr1_: Gαi2 and βarr1 couplings are lost (green). Bold outline indicates highly conserved residues/microswitches such as NPxxY, CWxP, etc. (**B**) Distribution of coupling profiles classes (see also scatter plot in **Figure S2**). (**C**) Emax distribution for CB_2_R mutants with a preferential Gαi2 coupling profile. (**D**) Distribution of coupling profiles for mutants in conserved regions. Percentages differ considerably from the ones of the full set of mutants in (**Figure 2B**). (**E**) Kernel density estimation of the distribution of conservation scores among PrefCoup_Gαi2_ mutants and Coup_Gαi2_βarr1_ mutants with a WT-like expression level (80% to 100%) showing a clear shift between both densities. A conservation score distribution for all coupling classes can be found in **Figure S4**.

To investigate whether a loss of βarr1 recruitment negatively impacts Gαi2 coupling, we plotted the Gαi efficacy (Emax_Gi_) distribution for mutants with impaired βarr1 recruitment (i.e., PrefCoup_Gαi2_ mutants) (**Figure 2C**). The plot reveals that PrefCoup_Gαi2_ mutants exhibit Gαi efficacies that are normally distributed around the WT level (**Figure 2C**), suggesting that abolishment of βarr1 recruitment does not necessarily affect the efficacy of G protein signaling. Of note, there is also a significant fraction of receptor mutants (12%) incapable of coupling to both Gαi and βarr1 (**Figure 2B**). These impactful mutations are enriched in highly conserved regions (e.g. microswitches, highlighted with a black circle, **Figure 2A**) and represent approximately 50% of highly conserved residues (**Figure 2D**).

To obtain a more comprehensive understanding of the implication of residue conservation for receptor functionality, we plotted its distribution for mutants with preserved Gαi2 and βarr1 coupling (Coup_Gαi2_βarr1_) compared to the ones with preferential Gαi2 coupling (PrefCoup_Gαi2_). To exclude a potential impact through differential receptor expression level, we only plot mutations with a WT-like expression level (i.e., 80% to 120%). We find that PrefCoup_Gαi2_ mutants (orange line, **Figure 2E**) show a clear shift toward higher conservation scores compared to Coup_Gαi2_βarr1_ mutants (blue line, **Figure 2E**). Our data highlight the relationship between the evolutionary conservation and the receptor coupling preference.

### Preferential Gαi2-coupled mutants are located in proximity to network connections with high information transmission

We hypothesize that the receptor dynamics and the stabilization of specific receptor states through an allosteric communication network (ACN) are crucial for the functional response of the CB_2_R. In this framework, PrefCoup_Gαi2_ mutants likely exert their effects by perturbing the allosteric communication network in the CB_2_R compared to mutants with preserved coupling to Gαi2 and βarr1 (Coup_Gαi2_βarr1_).

To test this hypothesis, we performed all-atom molecular dynamics (MD) simulations and sampled the ACN for the WT CB_2_R (WT ACN) in complex with the agonist HU-210 over 2 μs accumulated simulation time (5 x 400 ns). The ACN is computed based on contact frequencies including hydrophobic interactions, hydrogen bonds and water-mediated interactions. Then we identified the 100 shortest pathways on this network (LigACN) that mediate the communication from the orthosteric ligand binding site to the intracellular coupling site (**Figure 3A**). To estimate information transmission of a contact in the ACN, we computed the frequency of their appearance in 100 shortest pathways (see methods, **Figure 3A** and **B**). For our interpretation, we treat the LigACN as the principal contact network, which is stabilized by the ligand to promote a specific receptor state and functional response. In contrast, the entire WT ACN encompasses the complete allosteric connectivity within CB_2_R and provides valuable insights into connections that link remote receptor regions to the LigACN. These connections represent a resource that can be harnessed by allosteric modulators, including membrane lipids, small molecules, or other interacting proteins, to fine-tune receptor conformational states and, consequently, modulate its response.

**Figure 3.**
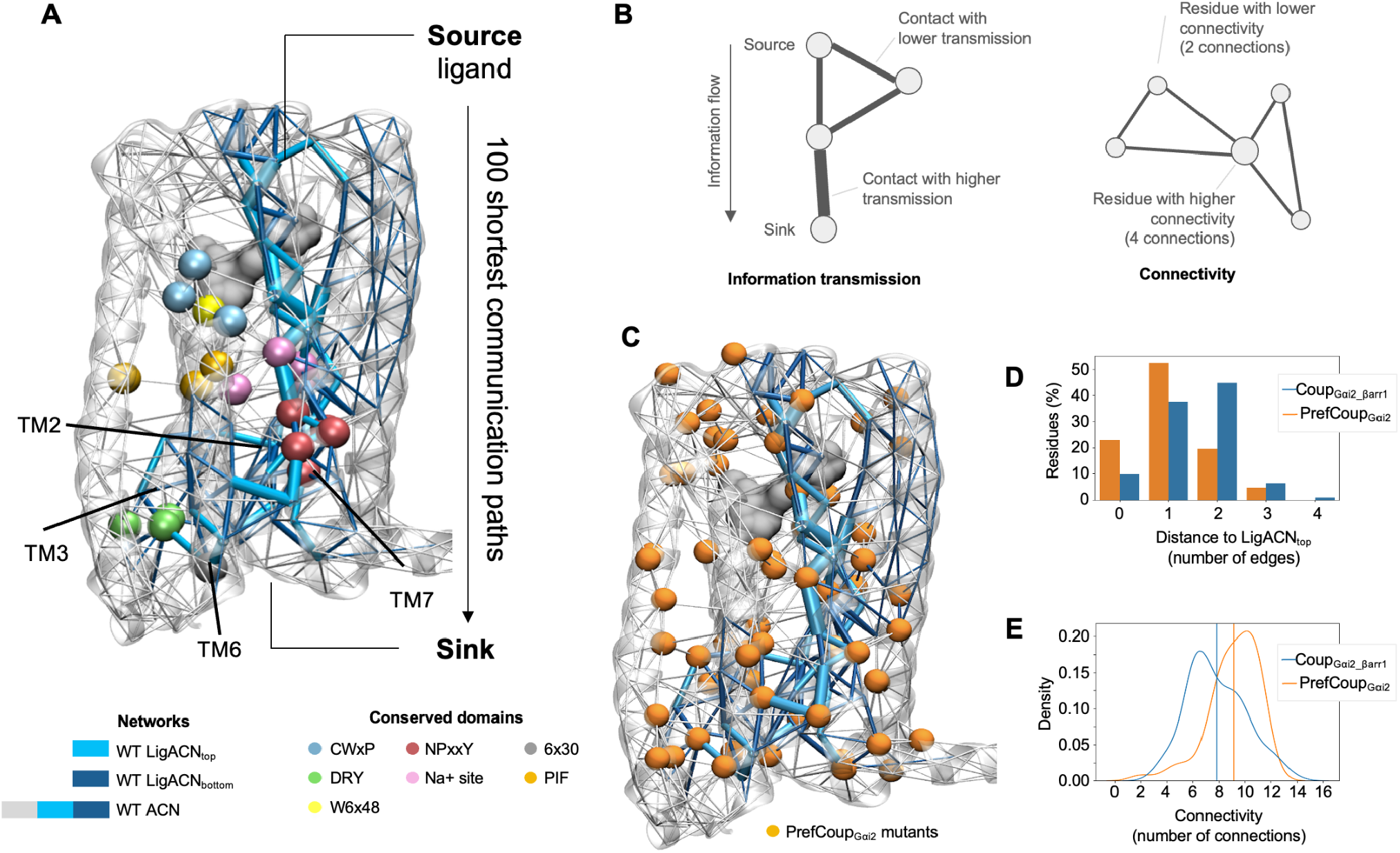
Relationship between PrefCoup_Gαi2_ mutants and the ACNs. (**A**) Depiction of the entire allosteric communication network in the WT CB_2_R (WT ACN) in relation to the highly conserved domains. The WT ACN includes the ligand-stabilized ACNs highlighting contacts with top (LigACN_top_) and bottom (LigACN_bottom_) contributions to the information transmission in the CB_2_R. The thickness of the contacts is proportional to their information transmission between the ligand and the intracellular receptor site (see methods). (**B**) Schematic illustration of information transmission and the connectivity of a specific node in the network. (**C**) PrefCoup_Gαi2_ mutants are shown as orange spheres on the WT ACN alongside the LigACN_top_ and LigACN_bottom_. (**C**) Distance of PrefCoup_Gαi2_ and Coup_Gαi2_βarr1_ mutants to LigACN_top_. The x-axis represents the number of edges from a mutation position (preferential Gαi2 coupling mutants in orange or mutants with preserved coupling in blue) to the closest residue from the LigACN_top_. The y-axis represents the percentage of mutants at a given distance, facilitating comparison between the coupling profiles. (**D**) Connectivity distribution plot of PrefCoup_Gαi2_ mutants (orange) and Coup_Gαi2_βarr1_ (blue). Connectivity for each residue position is computed as the number of connections in the network (see **Figure 3B**). Vertical lines indicate the mean value of each distribution.

A first inspection of the LigACN stabilized by our reference agonist HU-210 shows that it comprises several residues crucial for receptor function, known as conserved microswitches (**Figure 3A**). These microswitches include residue R131^3x50^ from the DRY motif, residue N291^7x45^ and S292^7x46^ in the sodium binding site, and all five residues in the NPxxY motif N295^7x49^ to Y299^7x53^ (the GPCRdb numbering scheme is used ^21^). The presence of these highly conserved residues within the LigACN strongly supports its relevance in transmitting information and maintaining the protein’s function. Furthermore, we find that the LigACN elicits a strong communication through the upper and lower part of TM7 as well as the lower parts of TM2, 6 and 3.

We then determined the distance of PrefCoup_Gαi2_ mutants to connections of high information transmission in the LigACN (LigACN_top_, see method section) (**Figure 3C**). Interestingly, we find that PrefCoup_Gαi2_ mutants are more populated closer to the LigACN_top_ (i.e. 0 to 1 connections, **Figure 3D**) whereas Coup_Gαi2_βarr1_ mutants at larger distances (2 to 4 edges). We verified that this closeness was statistically significant by comparing its mean value to a null distribution built using random sets of residues (p-value 0.006). On top of this proximity, inspection of the protein contact network revealed that PrefCoup_Gαi2_ mutants on average have higher connectivity (i.e., a residue has more connections to other residues) than Coup_Gαi2_βarr1_ mutants (**Figure 3B** and **D**). Both proximity to highly information-transmissive connections in the LigACN and the higher connectivity rationalize why these residues have a significant functional impact on the receptor’s coupling profile.

### Multiple molecular mechanisms initiate Gαi2 preferential coupling

To obtain a deeper mechanistic understanding of changes in the ACN that drive preferential Gαi2 coupling as a result of the loss of βarr1 recruitment, we carried out additional MD simulations of a set of 35 mutants from the 360 point mutations including 20 PrefCoup_Gαi2_ mutants and 15 Coup_Gαi2_βarr1_ (**Figure 4A, Supplementary Table S1**). The selection of mutants was done based on two criteria: (1) mutants show WT-like expression levels, as a reduced/increased receptor expression can significantly alter receptor signaling^22^ and (2) mutants are located in receptor regions that are structurally resolved. With the aim to understand the initial ACN re-arrangements for preferential Gαi2 coupling as a result of βarr1 loss, we started from an inactive receptor state (PDB ID 5ZTY, see method section). For each simulated mutant, we monitored the dynamics of formation and disruption for different interaction types within the entire receptor contact network. The interaction stability for thousands of interatomic connections was quantified by computing the contact frequency for the last 400 ns in 5 replicates (i.e. 2 μs total analysis time per mutant) (**Figure 4B**).

**Figure 4.**
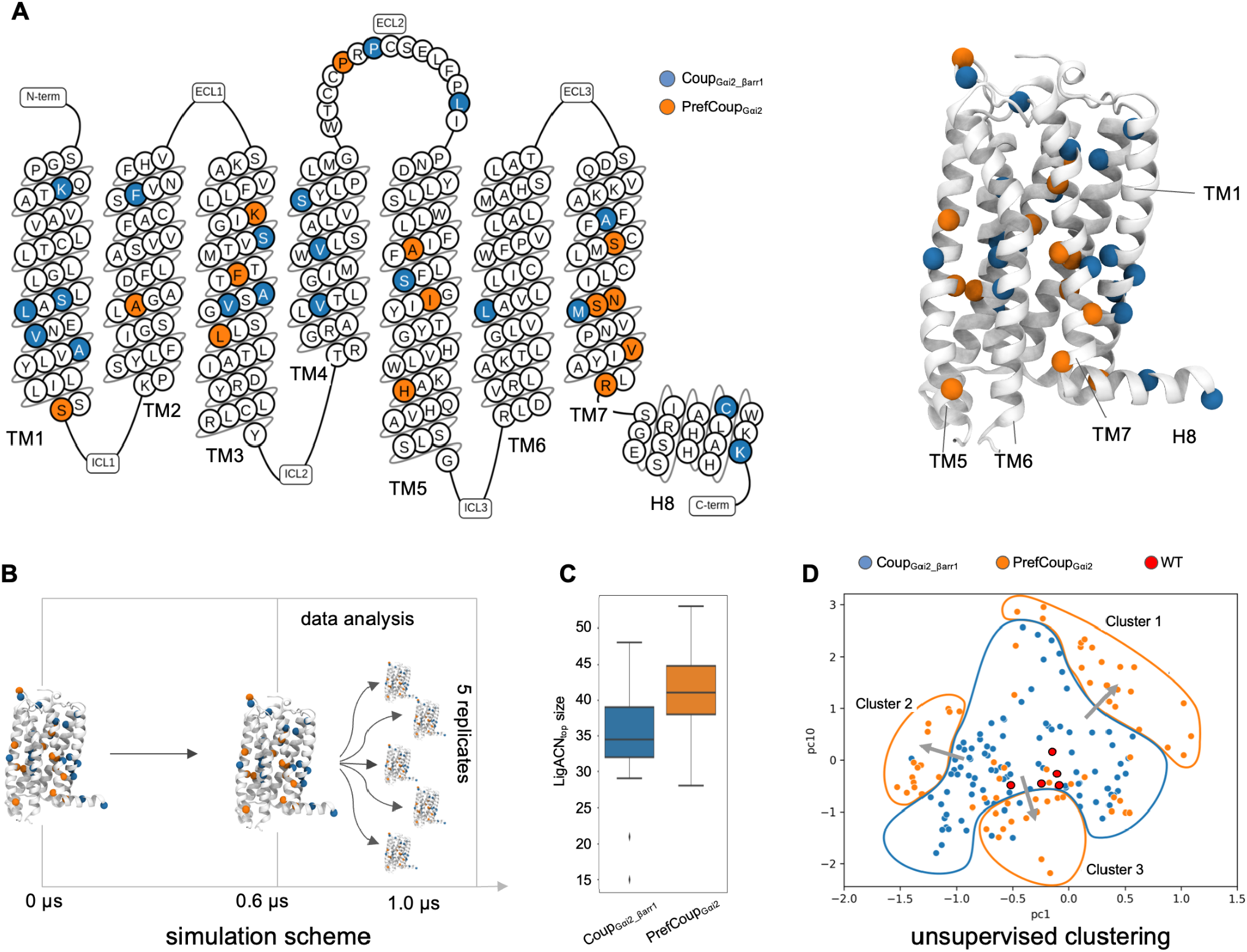
Clustering of simulated mutants based on alterations in the intramolecular contact network. (**A**) Location of simulated mutants and their coupling profiles shown on a snake plot (left) and the CB_2_R structure (right). PrefCoup_Gαi2_ mutants are shown in orange and Coup_Gαi2_βarr1_ in blue. (**B**) MD simulations scheme: the initial system was simulated in production conditions for 0.6 μs. After that, 5 replicates were respawned applying random perturbations to the atom speeds. These 5 replicates were evolved for further 0.4 μs reaching a total evolution time of 1 μs. This yields a total simulation time of 2.6 μs per mutant. The respawned replicates (5 x 0.4 μs) are used for data analysis (**C**) The network size of LigACNs (i.e., number of nodes) for PrefCoup_Gαi2_ mutants compared to WT and Coup_Gαi2_βarr1_ mutants. The data for LigACNs including interatomic contacts in the ACN for all 35 mutants is found in **Supplementary Data 2**. (**D**) Unsupervised clustering of MD data based on contact frequencies. Projection of the first (PC1) and tenth principal components (PC10) reveals clusters 1 to 3 for PrefCoup_Gαi2_ mutant systems (encircled with an orange line). The Coup_Gαi2_βarr1_ cluster (in blue) stands in the middle of the plane. The WT control simulations are highlighted in red.

First, we investigated if PrefCoup_Gαi2_ mutants induce alteration in the ligand-stabilized ACNs. In fact, we find that individual PrefCoup_Gαi2_ mutants typically induce larger LigACNs compared to Coup_Gαi2_βarr1_ mutants, i.e. increase the number of nodes in the pathways from the ligand to the intracellular receptor site (**Figure 4C**). It seems that Coup_Gαi2_βarr1_ mutants disrupt information transmission pipelines in the WT LigACN. This disruption leads to a branching of the information flow along longer pathways including more residue nodes, likely resulting in a less efficient transmission of information. The functional readout of those mutants suggests that the inclusion of more residues in the communication pipeline and in turn less efficient information transmission is linked to a loss of β-arrestin recruitment in the CB_2_R.

To extract further structural information relevant for preferential CB_2_R coupling, we reduced the dimensionality of simulated contact networks by applying principal component analysis (PCA). Interestingly, projecting the data on a principal component (PC) plane defined by PC1 and PC10 reveals a separation of Coup_Gαi2_βarr1_ mutants (blue points) and the PrefCoup_Gαi2_ mutants (orange points) (**Figure 4C**). In other words, preferential Gαi2 coupling and loss of βarr1 loss recruitment in our dataset is described by one PC (PC1) with high explained variability (13%) and a second PC (PC10) with less explained variability (3%). Apparently, there are also contacts with high variability (i.e., PC2 to 9) that are not related to preferential Gαi2 coupling in the CB_2_R and that could be linked to other receptor functions.

Interestingly, the spreading of the mutants that preserve coupling (Coup_Gαi2_βarr1_, blue points) around the true WT system (red points) indicates that some variability in the contact network is tolerated for preserving coupling with an Emax > 50%. Intriguingly, we find that PrefCoup_Gαi2_ mutants (orange points) tend to shift further away from the true WT center (red points) (**Figure 4D**). Apparently, these mutations perturb specific elements (i.e., stabilization or destabilization) in the contact networks that fall outside the tolerated region of the Coup_Gαi2_βarr1_ mutants (blue points). In addition, we find that PrefCoup_Gαi2_ mutants move in three different directions forming clusters 1 to 3. Hence, each cluster appears to differently disrupt the contact network, leading to impaired βarr1 recruitment and preferential Gαi2 coupling. Ultimately, this suggests that different mechanisms are implicated in preferential Gαi2 coupling and the loss of βarr1 recruitment in the CB_2_R. This could also explain why some PrefCoup_Gαi2_ mutants fall within the blue cluster of preserved coupling as they likely exploit another molecular mechanism that is not projected into the plane of PC1 and PC10.

Furthermore, we observe that the three PrefCoup_Gαi2_ clusters (orange points) are located at different distances from the true WT simulations (red points). Specifically, cluster 3 is very close to the true WT simulations compared to clusters 1 and 2. This suggests that cluster 3 - specific contact stabilities are more sensitive and smaller network alterations can push the CB_2_R from a preserved coupling profile to a PrefCoup_Gαi2_ profile.

Of note, we also simulated the same 35 mutants including 15 Coup_Gαi2_βarr1_ and 20 PrefCoup_Gαi2_ mutants in the active CB_2_R structure coupled to the Galpha subunit (PDB ID 6KPF). Interestingly, we were not able to obtain conclusive results discriminating PrefCoup_Gαi2_ from the Coup_Gαi2_βarr1_ profile. This is likely because binding of the Gα subunit to the intracellular coupling site of the CB_2_R stabilizes the ACN which hampers the analysis.

### Contact alterations in conserved molecular switches are involved in Gαi2 preferential coupling

To capture in more detail the structural and dynamic alterations in the receptor contact network that lead to Gαi2 preferential coupling, we fitted a logistic regression for each PrefCoup_Gαi2_ cluster to separate them from the Coup_Gαi2_βarr1_ group (see method). Top interaction features (orange color scaled connections) that distinguish preferential Gαi2 from the preserved coupling profile are plotted on the allosteric communication network (WT ACN, gray) including the ligand-stabilized ACN (cyan) (**Figure 5A**). The obtained representation indicates that mutational perturbation in preferential Gαi2 coupling for clusters 1 to 3 (orange color scale) involves a complex and multifaceted modulation of the WT ACN (gray plus cyan) and the corresponding LigACN (cyan). To further simplify the interpretation of this complex perturbations of the WT ACN, we focus on highly conserved receptor regions with known relevance for receptor function such as the CWxP motif, allosteric sodium binding site, NPxxY motif, and the DRY motif. Cluster-specific modulations are depicted in **Figure 5B-D** (detailed structural description can be found in **Supplemtenary Note 1**). A cluster summary (**Figure 5E**) reveals that the sodium binding site and the DRY motif are modulated in all three identified clusters, the CWxP only in clusters 1 and 2, and the NPxxY motif only in cluster 3. Based on the finding that the DRY motif and the sodium binding site are implicated in all three clusters, it is tempting to speculate that they play the most important role in preserving CB_2_R coupling to Gαi2 and βarr1 upon mutational perturbations. Another interesting observation is that detected perturbations around the sodium binding site, CWxP and DRY motifs involve contact (de)stabilization effects through different residues (**Figure 5E**). This corroborates the finding that multiple molecular mechanisms can converge into the same receptor response, i.e. loss of βarr1 recruitment while preserving Gαi2 coupling with an Emax > 50%.

**Figure 5.**
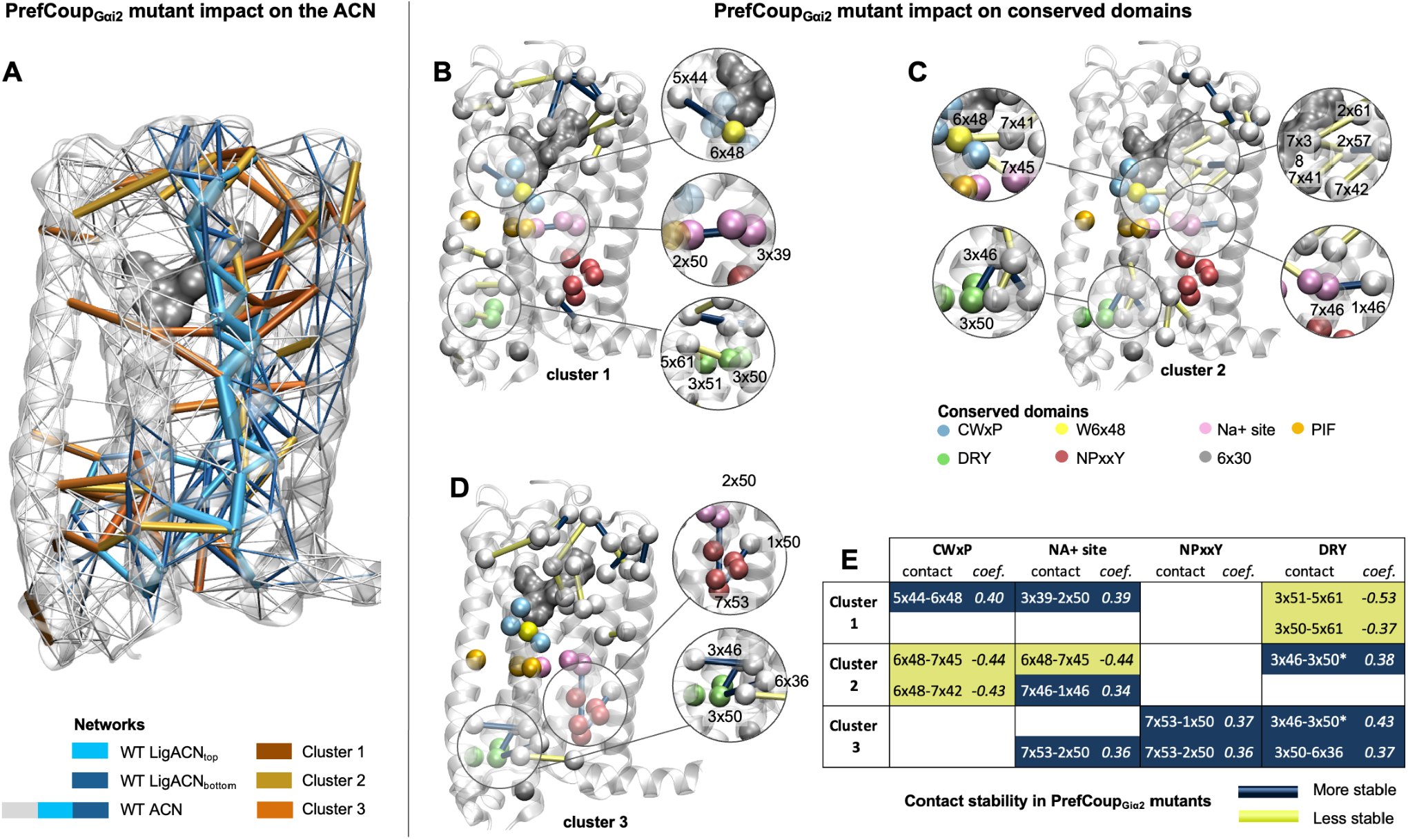
Characterization of PrefCoup_Gαi2_ mutant clusters. (**A**) Perturbation map of the ACN induced by PrefCoup_Gαi2_ mutants. Top interaction features for PrefCoup_Gαi2_ cluster 1 (dark brown), 2 (light brown) and 3 (orange). The interactions are plotted together with the computed ACNs for the WT CB_2_R. For instance, such depiction highlights allosteric connections (brown to orange) to the ligand-stabilized ACN (cyan) that are perturbed in preferential Gαi2 coupling. (**B-D**) Mutant impact on highly conserved receptor regions for clusters 1 to 3. Non-conserved residues which are part of the modulation are represented as white spheres whereas the corresponding contact is highlighted as a cylinder indicating the change of contact stability compared to conserved coupling (Coup_Gαi2_βarr1_): navy blue corresponds to increased interaction stability in the PrefCoup_Gαi2_ mutants, yellow indicates decreased interaction stability. The thickness of the cylinders is proportional to the absolute value of the coefficient for that interaction in the logistic regression model. (**E**) Summary of mutant-perturbed domains of high conservation (i.e., CWxP, Na+ site, NPxxY, DRY) for PrefCoup_Gαi2_ mutant clusters 1 to 3. For each cell, we indicate the implicated residues in the contact and logistic regression coefficient. Each cell is colored according to the sign of the logistic regression coefficient (navy blue for positive and yellow for negative coefficients).

## Discussion

Allosteric communication networks are highly complex dynamic entities that drive receptor conformational states and in turn determine their final functional outcome as a response to environmental changes (e.g., mutations, ligand stimulation). Here, we investigated the dynamics of such networks and their modulation related to preferential signaling of the CB_2_R by integrating functional readouts from an alanine/valanine scanning mutagenesis library with all-atom computer simulations. The results obtained from our study shed light on several key aspects that link receptor structure and dynamics to function.

First of all, our functional readouts suggest that Gαi2 coupling is more robust to mutational perturbations compared to βarr1 recruitment in the CB_2_R (**Figure 2B**). Our observation is in line with a similar study on β_2_ adrenergic receptor (β_2_AR) where more than 75% of mutational substitutions retain the receptor’s G protein coupling profile^23^. However, it is important to mention that this is not a general tendency for GPCRs as shown recently for the angiotensin II type 1 receptor (AT1R)^11^. For this receptor, β-arrestin recruitment is more resistant to mutations compared to G protein binding. This could possibly be related to the intrinsic propensity of GPCRs to differentially engage in interactions with β-arrestins. While class A GPCRs for recruitment (such and β_2_AR and CB_2_R) interact with arrestins only transiently, class B receptors (V2R and AT1R) form more stable interactions which are potentially more resistant to disruptions by single-point mutagenesis^24^.

Plotting the Gαi2 preferential mutants on the static CB_2_R structure helps obtain a first impression of their potential impact based on their location (**Figure 2**). However, such representation falls short when studying the molecular mechanism that drives the modulation of a receptor that is in constant motion. Here, we demonstrate that the intrinsic CB_2_R flexibility can be captured with atomistic resolution through allosteric communication networks within the receptor in which each interatomic connection is characterized through a distinct contact frequency. Further network processing using graph theory highlights how the agonist HU-210 allosterically communicates with the intracellular coupling site of the CB_2_R mainly through TM7 (**Figure 3A**). The relevance of TM7 related agonist binding goes along with a previous study for the β_2_ adrenergic receptor despite using a different approach. Applying the concept of correlated movements for the detection of allosteric communication pipelines ^25^, the authors find that the inverse agonist-bound inactive states uses mainly TM6 whereas agonists-bound active states favor TM7 as the main communication pipeline.

Investigating the mutant position, their information transmission and connectivity within such networks reveals a complex modulation of CB_2_R coupling (**Figure 3B**). For a more detailed description, we introduced the concept of the ligand-stabilized ACN (LigACN) which is part of the overall CB_2_R ACN. The LigACN is important for stabilizing the receptor in a specific conformational ensemble and therefore directing the functional outcome of the ligand-stimulated receptor. Importantly, we find that Gαi2 preferential mutants are located closer to connections of high transmission in the ligand-stabilized ACN (**Figure 3B-C**) and are of higher network connectivity (**Figure 3D**). From this, we can conclude that both information transmission and connectivity are critical residue properties for modulating the allosteric network in the CB_2_R and in turn its ability to recruit βarr1. Mechanistically, this can be explained as follows: mutant perturbations of allosteric connections (i.e., loosening or strengthening) in the WT ACN and specifically in the LigACN shift the conformational CB_2_R ensemble from states with favorable affinities for multiple transducer proteins (e.g. Gαi, βarr1, others) to states that lose βarr1 while preferentially preserving Gαi2 with Emax > 50% (**Figure S3**).

Our findings are in line with previous studies for GPCRs^16,26^ and other protein classes^27^ showing that disruption of allosteric communication networks through mutations results in a significant impact on protein function. This also has implications for allosteric modulators that can fine-tune function by perturbing allosteric connections between different protein regions^28^. However, it is worth noting that not all mutations occurring within our computed LigACN consistently exhibit the same level of efficiency in altering Gαi2 preferential coupling. This is not surprising as those positions might be involved in other functional consequences (e.g., coupling to other (un)known signaling partners) that are not taken into account in our study. Further investigations are required to better understand these functional relationships.

Ultimately, the most important finding of our study is related to the observation that there are multiple molecular mechanisms that can induce the same functional response, i.e. preferential Gαi2 coupling. In fact, this goes along with the applied concept of allosteric communication networks that includes regulatory triggers from different receptor regions. Principal component analysis (**Figure 3D**) and logistic regression (**Figure 4**) pinpoint three separate mechanisms that regulate highly conserved receptor motifs such as CWxP, sodium binding site, NPxxY and DRY. The involvement of those microswitches in impairing β-arrestin recruitment has been reported in different individual studies (i.e., DRY motif^28–31^, NPxxY motif^32^, the toggle switch ^33^, and the sodium binding site ^34^). Importantly, we find that the same molecular switch can be perturbed through different combinations of interactions and contact stabilities as highlighted by our PrefCoup_Gαi2_ clusters 1 to 3 (**Figure 4B-D**). Ultimately, we anticipate the existence of other molecular mechanisms based on the observation that not all PrefCoup_Gαi2_ mutants in our dataset (e.g., S61A on the bottom of TM1) fall into one of the three PrefCoup_Gαi2_ clusters, but are located in the preserved signaling cluster (**Figure 3C**). Moreover, we expect that additional mechanisms will come into play when changing experimental conditions such as the choice of agonist or the type of mutational substitution (e.g., as occurring in single nucleotide polymorphisms). Given the multitude of molecular mechanisms that induce preferential Gαi2 coupling suggests that biased ligands may not all operate through a shared universal mechanism. Instead, it is possible that different classes of ligands achieve bias by employing distinct structural mechanisms. By delving into these unique mechanisms, we can advance our understanding of biased signaling, target GPCR-mediated signaling pathways more effectively, and eventually broaden our repertoire of therapeutic approaches.

In conclusion, our study provides unprecedented insights into the intricate interplay of multiple triggers on complex allosteric communication networks in the CB_2_R and their final functional outcomes. Potent triggers of such changes in ACNs that lead to distinct intracellular signaling responses are characterized by high connectivity and their close vicinity to high information-transmissive connections. These findings have important implications for elaborating drug design strategies aimed to tailor receptor response towards more efficacious and safer drugs. However, given the complexity of nature and the involvement of multiple molecular mechanisms, further research is necessary to fully comprehend these intricate processes and their potential applications in drug development.

## Methods

### Mutant generation

#### Plasmids and mutagenesis

Human cannabinoid CB_2_ receptor gene was optimized for expression in mammalian systems and synthesized (Genewiz). The CB_2_R gene was fused to an N-terminal signal sequence, SNAP-tag, TwinStrep-tag and a C-terminal 1D4-tag and cloned into pcDNA4/TO vector. Single-point mutants were generated for all 360 CB_2_ residues where non-alanine amino acids were replaced by alanine and alanines were replaced by valines. For that purpose, PCR with mutation-containing primers was used as described in Heydenreich et al.^35^. Plasmids encoding β-arrestin1-RlucIl^36^, human GRK2^37^, rGFP-CAAX^17,37^, and wild-type Gαi2 protein and an effector protein fused with RlucII (Rap1GAP-RlucII)^20^ were previously described. Briefly, for the G protein activation assay, receptor and wild-type Gαi2 subunit were co-transfected with Rap1GAP-RlucII effector and rGFP-CAAX, while in β-arrestin1 recruitment assay, receptor was co-transfected with β-arrestin1-RlucII, wild-type GRK2 and rGFP-CAAX. In both assays an increase in BRET signal was monitored upon ligand-induced Rap1GAP or β-arrestin recruitment to the plasma membrane.

### Signaling profile determination

#### Transfection

Human embryonic kidney (HEK) 293T cells^38^ were cultured in DMEM, 4.5 g/L glucose, with L-glutamine and phenol red, without sodium pyruvate (Wisent Inc., Montreal, Canada), supplemented with 10% Newborn Calf Serum (NCS, iron fortified, Wisent Inc., Montreal, Canada) and 1x Penicillin-Streptomycin (PS, Wisent Inc., Montreal, Canada) in adherent culture at 37°C, 5% CO_2_. For transfection, medium without phenol red was used. Cells were dissociated using 0.05% Trypsin with 0.53 mM EDTA (Wisent Inc., Montreal, Canada) and transiently transfected with receptor and biosensor DNA using 25 kDa linear polyethylenimine (Polysciences Inc., Warrington, PA, US) in Cellstar® PS 96-well cell culture plates (Greiner Bio-One, Germany) at a density of 20 000 cells per well.

### BRET experiments

Coelenterazine 400a (Prolume Ltd., Pinetop, AZ, US) was used as the luciferase substrate. HU-210 was purchased from Toronto Research Chemicals (Toronto, ON, Canada) BRET experiments were performed 2 days post-transfection, when culture medium was replaced by 100 μL Tyrode’s buffer (137 mM NaCl, 1 mM MgCl_2_, 1 mM CaCl, 0.9 mM KCl, 3.6 mM NaH, 11.9 mM NaHCO3, 25 mM HEPES, 5.55 mM D-glucose, pH = 7.4). A volume of 0.4 μL of ligand was added to cell plates using a BiomekFX Lab Automation Workstation (Beckman) and a previously calibrated pin tool, followed by incubation at 37°C for 5 minutes in a Cytomat 6001 (Thermo Scientific). Coelenterazine 400a working solution (50 μM DBC and 1% Pluronic F-127 in Tyrode’s buffer) was added with a Multidrop Combi (Thermo Scientific) to a final concentration of 5 μM and shaken for 10 seconds. The plates were further incubated for 5 minutes at room temperature before the BRET signal was read on a Synergy Neo (Biotek) equipped with dual photomultiplier tubes (PMTs) (emission recorded at 410 and 515 nm, gain settings of 150 for each PMT and 1.2 s integration time).

### Cell surface ELISA measurements

HEK293T cells were transfected as above using only receptor-coding plasmids in poly-D-lysine coated 96-well plates and incubated at 37°C and 5% CO_2_ for 2 days. Cells were washed with PBS (200 μL/well) and fixed with 3% paraformaldehyde (50 μL/well) for 10 min, followed by three washing steps (100 μL/well, PBS + 0.5% BSA). Receptor was detected by primary rabbit anti-SNAP antibody (50 μL/well of 1:2000 dilution, GenScript, 1 h at RT) and anti-rabbit HRP antibody (50 μL/well of a 1:1000 dilution, GE Healthcare, 1h at RT), with washing steps as above, followed by three wash steps with PBS. SigmaFast solution (100 μL/well) was added, followed by incubation at RT protected from light, until color change was observed. To stop the reaction, 25 μL 3M HCl were added to each well and 100 μL of the solution were transferred to a new transparent 96-well plate for the measurement. The absorbance at 492 nm was measured using a Tecan GENios Plus microplate reader, reporting on the amount of receptor expressed.

### Effect of the receptor expression level on maximal ligand-induced response

Receptor titrations were performed to determine the effect of receptor expression level on Emax values of both pathways. Various amounts of CB_2_ DNA were transfected (0, 0.1, 0.5, 1, 2.5, 5, 10, 25, 50, 100, 150, 200% of the amount used for each biosensor), together with other biosensor components in duplicates. Two days post-transfection one replicate was used to perform

ELISA, while the other replicate was used to measure BRET signal upon stimulation with 1 μM HU-210, as described above. Change in BRET signal (Emax) vs. amount of CB_2_R transfected (percentage of the amount used for each biosensor), and cell-surface expression level (determined by ELISA) vs. amount of CB_2_R transfected were measured. Correlations of the Emax and ELISA-measured expression levels were determined for each pathway, and data fit equation parameters (linear regression for Gαi2 activation and non-linear hyperbolic fit for β-arrestin1 recruitment) were used to correct Emax output of each mutant for its previously measured expression level (**Supplementary Data 1**).

### BRET data analysis

The BRET response was determined by the ratio of the light intensity emitted by the acceptor (rGFP, measured at 515 nm), over the light intensity emitted by the BRET donor (RlucII, measured at 410 nm). Single-replicate concentration-response curves were fitted to a Hill equation with a Hill coefficient of 1. For each concentration-response curve, the maximum response of signal (Emax) and the ligand concentration at half-maximal signal response (as pEC50) was determined using the custom software DataFitter (https://github.com/dbv123w/DataFitter), followed by averaging over biological triplicates. For further analysis, Emax of CB_2_R mutants was corrected for the mutant expression levels and normalized to the wt value (**Figure S6**), while pEC50 of each mutant was subtracted from the wt value. Both raw and corrected values are reported (**Supplementary Data 1**).

### Model generation and molecular dynamics simulation

As a starting point, we have selected the inactive crystal structure of the CB_2_R bound to the antagonist AM10257 (PDB code: 5ZTY^39^). The sequence of the receptor was reverted to the canonical human sequence using data from UNIPROT [P34972]. Residues between positions 222 and 235 of the ICL3 were removed and endings were capped using ACE and CT3 terminal patches. The initial position of the ligand was generated based on the position of the highly similar agonist - AM12033 co-solved in the active CB_2_R CryoEM structure (PDB ID 6KPC) (**Figure S5**)^40^. The resulting complex was embedded in a 54% POPC, 36% POPG and 10% cholesterol membrane, based on the coordinates available in the OPM database^41^. The pre-equilibrated membrane was generated in CHARMM-GUI^42^. Protonation states at a pH of 7.4 were assigned using Propka, and residues facing the membrane were manually curated to avoid charges facing the membrane. The obtained system was solvated with TIP3 waters, and the ionic strength of the solution was maintained at 0.15 using NaCl ions.

The obtained complex was equilibrated for 100 ns in NPT conditions, with constraints applied to the protein backbone and using the Berendsen barostat^43^. From the equilibrated system, we generated systems for each of the studied mutants (See list in **Supplementary Table S1**). Every system underwent a short 5 ns NPT equilibration phase to adjust the environment to the introduced mutation. Afterward, we applied a simulation scheme in which each mutant evolved for 1 μs simulation time in an NVT ensemble (**Figure 3B**). The last 400 ns were carried out in 5 replicates reaching a total simulation time of 2.6 μs per mutant (i.e, 600 ns plus 5* 400 ns = 2.6 μs).

All simulations were carried out using the ACEMD engine^44^. Parameters were obtained from the Charmm36M forcefield generated in November 2017^45^, with ligand parameters generated in Paramchem^46,47^ with the CgenFF forcefield^48,49^. The temperature was maintained at 310 K. For NPT runs we have employed a 2 fs timestep, and for NVT runs a 4 fs one. This was possible due to the hydrogen mass repartitioning scheme employed within ACEMD.

### Conservation metrics computation

To calculate the evolutionary conservation score for each residue of the CB_2_R, we obtained the structural sequence alignment of class A GPCRs from GPCRdb and input it into Capra et al.’s online tool with the Jensen Shannon Divergence (JSD) scoring method^18^. The default values were used for all parameters, including a window size of 3 and BLOSUM 62 matrices for both the background and target. The JSD measures the similarity between two probability distributions by comparing the distribution of amino acids from a sample in a particular position with the distribution of a set without evolutionary pressure^18^. The JSD ranges from 0 to 1, where 0 indicates identical distributions, and 1 means that the distributions have no overlap. Hence, a JSD score of 1 indicates that the two probability distributions are entirely dissimilar with no common features. We then used conservation scores to investigate the relationship between the impact of mutations on coupling affinities and the evolutionary pressure on each residue. To achieve this, we calculated the ratio of Gαi2 preferential mutants above each JSD score obtained in the analysis. For example, we found that out of all the residues with a conservation score of 0.36 or higher, 70% are Gαi2 preferential, compared to the 40% found in the whole sequence.

### Contacts computation

The contact computation in the simulations was done using the GetContacts package [https://getcontacts.github.io/]. All the interaction types supported by the software were used to compute interaction frequencies. Namely, hydrogen bonds, salt bridges, pi-cation, pi-stacking, t-stacking, hydrophobic, and van der Waals. Each of the interaction types has a set of conditions regarding the distance and relative angle of the two groups involved. The complete specifications of these criteria can be found in the GetContacts repository. After computing the interactions happening in each frame, the overall contact frequency between pairs of residues was computed. This was achieved by counting all the frames where any contact type was established between two residues and dividing it by the total number of frames of the simulation.

### ACN computation

*WT ACN computation*. To construct the allosteric network, we initially calculated the inter-residue contact stability network. This involved measuring the frequency of weak interactions between residue pairs across multiple simulation frames. From this network, we removed two types of interactions: (1) consecutive residues in the sequence, as their interactions were consistently established and provided limited useful information, and (2) interactions with very low frequencies that could occur by chance in only a few frames of the simulation.

*Ligand-stabilized ACN (LigACN)*. Using the resulting network, we determined the 100 shortest pathways between the ligand and each of the four selected residues in the intracellular domain for each replica of the simulation (**Figure 3B**). These specific residues, namely Arg131 (3x50), Asp240(6x30), Ser303 (8x47), Ser69 (2x39), were chosen based on their proximity to the G-protein in the bound conformation. Subsequently, we computed the information transmission for each edge in the network by computing their degeneracy. The edge degeneracy is calculated as the frequency of each edge in the suboptimal paths. To reduce noise, we applied a threshold to the degeneracy values, filtering out less significant edges from the allosteric communication pathway (**Figure 3A**).

The AlloViz v0.1 Python package (github.com/frannerin/AlloViz) was used to compute the allosteric communication network. The raw network was calculated using the GetContacts method, considering all salt bridges, pi-cation, aromatic, and hydrogen bond (direct and water-mediated) interactions in the trajectories. Following the protocol by Foutch et al. (2021), the raw interaction frequency networks were filtered to remove edges connecting neighboring residues in the protein sequence ^50^. Additionally, we defined a lower frequency cutoff of 0.1 to remove the noise generated by the random fluctuation of the system.

The 100 shortest (suboptimal) paths connecting the ligand (CHEMBL5085420) to each of the four selected residues were computed in each replica using Dijkstra’s algorithm as implemented in NetworkX [https://www.osti.gov/biblio/960616]. This resulted in a total of 2000 suboptimal paths.

### Distance to LigACN computation

We define the distance to the LigACN as the number of edges in the shortest path between the mutant residue of interest and any residue from the LigACN. To remove noise from the LigACN, we also wanted to filter out edges with low degeneracy from the LigACN. To achieve this, we computed the distance filtering out low degeneracy edges at various thresholds. We selected the threshold that maximized the difference between the PrefCoup_Gαi2_ mutants and a random distribution. This random distribution was built by randomly selecting 1000 sets of residues with the same size as the PrefCoup_Gαi2_ set and computing their mean distance to the LigACN. This selected threshold was used to define two sets of edges within the LigACN: the edges with low degeneracy (LigACN_bottom_) and the edges with higher degeneracy (LigACN_top_). Finally, we tested if the proximity of PrefCoup_Gαi2_ mutants to the LigACN_top_ was higher than could be expected by chance. For this, we calculate the quantile of the distance of PrefCoup_Gαi2_ mutants in the distribution of the distance of random residue sets.

### Definition of a balanced set of CB_2_R alanine mutants for unveiling the determinants Gαi2 preferential signaling

We compiled a balanced dataset of Coup_Gαi2_βarr1_ and Gαi2 preferential mutants according to the following criteria: (i) Coup_Gαi2_βarr1_ with Emax_Gαi2_ and Emax_βarr1_ > 50%, (ii) preferential Gαi2 coupling with an Emax_Gαi2_ > 50% and no measurable arrestin activity (**Supplementary Data 1, Figure S2**). As cell-surface expression can alter receptor signaling^22^, we prioritized the selection of mutants with WT-like expression levels. A final requirement for mutant selection was that mutants are in receptor regions that had been structurally resolved previously to allow their investigations in reliable three-dimensional dynamics models. The final set consists of 35 mutants of which 20 were Coup_Gαi2_βarr1_ and 15 were Gαi2 preferring mutants. The location of selected mutants in the CB_2_R is indicated in **Figure 4A**. Despite the mutant class definition relying exclusively on Emax, it is not ignoring EC50. This is because all the mutants without measurable efficacy also presented a potency of 0. So, independently of which criteria, Emax or EC50, is used, the compositions of the groups would be identical.

### Cluster definition

To identify differences in the contact patterns between the Gαi2 preferential and Coup_Gαi2_βarr1_ mutants first we need to identify sets of Gαi2 preferential mutants with similar contact patterns. To do so, first, we reduced the dimensionality of the dataset by projecting the contact stabilities into the first 10 principal components. Then, we visually explored all 45 pairwise combinations of such components. This inspection revealed the presence of a set of outlier simulations, all of them belonging to the mutant <outlier mutant number>. After the removal of this mutant, we noticed a strong spatial dependence of the coupling profile on the plane defined by principal components 1 and 10. We manually assigned Gαi2 preferential mutants in three clusters and Coup_Gαi2_βarr1_ mutants in one. This classification was based spatial disposition of the mutants.

### Cluster characterization

To characterize each cluster, we used logistic regression to differentiate between the wild type and the biased cluster. Because the model couldn’t generalize well with divided training and validation data, we opted to train a regularized model with high bias on the entire dataset. After training, we extracted the coefficients of each predictor variable, sorted them based on absolute value, and selected the top 20 variables. We then associated the coefficient with the residues involved in the interaction that formed the predictor variable. Model and model training was done using the scikit-learn Python module. From all interactions that were highlighted by the highest coefficients in absolute value from the logistic regression we paid special attention to those involving conserved residues and interactions that were densely packed in a specific region of the receptor.

## Acknowledgments

AMP acknowledges support from the Instituto de Salud Carlos III FEDER (FI19/00037). JS acknowledges funding from the Instituto de Salud Carlos III (ISCIII) (AC18/00030) and the Instituto de Salud Carlos III (ISCIII) & co-funded by the European Union (PI18/00094). This work was supported by the Swiss National Science Foundation grants 135754 and 159748 and SNF Sinergia CRSII3_141898 to DBV. TM was awarded the Swiss National Science Foundation Doc.Mobility supplement and the EMBO Short-term Fellowship (420-2016). AMP, TM, TMS, VLM, FMH, DBV, JS are members of COST Action CA18133 “ERNEST”.

## Author contributions

TM carried out the alanine scan for the CB_2_R under the supervision of DBV and MB and the signalling assays under the supervision of MB and DBV. FMH, BP and JD contributed to data collection. CLG designed some of the biosensors used in this study. TF contributed to the initial data analysis. WG, ACR and UG participated in ligand selection for the alanine scanning signalling screen and provided unique CB_2_-selective compounds. DBV wrote the DataFitter software for high-throughput data analysis. AMP, VLM and TMS carried out the molecular dynamics simulations and data analysis under the supervision of JS. AMP and JS wrote the manuscript. TM and DBV contributed to the manuscript writing. All authors revised the manuscript. DBV and JS supervised and coordinated the whole project.

## Competing interests

MB is the president of the scientific advisory board of Domain Therapeutics, a biotech company to which biosensors used in this study were licensed for commercial use. DBV is founder of Z7 Biotech Ltd, an early-stage drug discovery contract research organization. All other authors declare no conflict of interest.

## Data and materials availability

Data supporting the findings of this manuscript are available as a **Supplementary Information** and **Supplementary Data 1 to 2**. MD simulations are deposited at the GPCRmd database (www.gpcrmd.org). Additional data supporting the findings are available from the corresponding authors upon reasonable request.

## Supporting file list

Supplementary Information: Contains a supplementary note describing in detail structural features of cluster 1 to 3, supplementary figures and tables.

Supplementary Data 1: Characterization of alanine/valine mutants of the CB_2_R including expression levels, Emax, and pEC50 for Gαi2 and βarr1 and classified coupling profile.

Supplementary Data 2: Transmission information computed as degeneracy for 35 simulated mutants.

## Abbreviations

CB_2_R: Cannabinoid 2 receptor
Gαi: inhibitory G alpha subunit
βarr1: β-arrestin 1
GPCR: G protein-coupled receptor
ACN: Allosteric communication network
LigACN: Ligand-stabilized ACN
MD: Molecular dynamics
TM: Transmembrane domain

## Notes

### Competing Interest Statement

The authors have declared no competing interest.

## References

1. Rosenbaum, D. M., Rasmussen, S. G. F. & Kobilka, B. K. The structure and function of G-protein-coupled receptors. Nature 459, 356–363 (2009).

2. Maresz, K. et al. Direct suppression of CNS autoimmune inflammation via the cannabinoid receptor CB1 on neurons and CB_2_ on autoreactive T cells. Nat. Med. 13, 492–497 (2007).

3. Ashton, J. C. & Glass, M. The cannabinoid CB_2_ receptor as a target for inflammation-dependent neurodegeneration. Curr. Neuropharmacol. 5, 73–80 (2007).

4. Maccarrone, M. et al. Goods and Bads of the Endocannabinoid System as a Therapeutic Target: Lessons Learned after 30 Years. Pharmacol. Rev. 75, 885–958 (2023).

5. Pacher, P., Steffens, S., Haskó, G., Schindler, T. H. & Kunos, G. Cardiovascular effects of marijuana and synthetic cannabinoids: the good, the bad, and the ugly. Nat. Rev. Cardiol. 15, 151–166 (2018).

6. Di Marzo, V. Targeting the endocannabinoid system: to enhance or reduce? Nat. Rev. Drug Discov. 7, 438–455 (2008).

7. Smith, J. S., Lefkowitz, R. J. & Rajagopal, S. Biased signalling: from simple switches to allosteric microprocessors. Nat. Rev. Drug Discov. 17, 243–260 (2018).

8. Luttrell, L. M. Minireview: More than just a hammer: ligand ‘bias’ and pharmaceutical discovery. Mol. Endocrinol. 28, 281–294 (2014).

9. Sivertsen, B., Holliday, N., Madsen, A. N. & Holst, B. Functionally biased signalling properties of 7TM receptors - opportunities for drug development for the ghrelin receptor. Br. J. Pharmacol. 170, 1349–1362 (2013).

10. Morales-Pastor, A. et al. In Silico Study of Allosteric Communication Networks in GPCR Signaling Bias. Int. J. Mol. Sci. 23, 7809 (2022).

11. Cao, Y. et al. Unraveling allostery within the angiotensin II type 1 receptor for Gα and β-arrestin coupling. Sci. Signal. 16, eadf2173 (2023).

12. Torrens-Fontanals, M., Stepniewski, T. M., Gloriam, D. E. & Selent, J. Structural dynamics bridge the gap between the genetic and functional levels of GPCRs. Curr. Opin. Struct. Biol. 69, 150–159 (2021).

13. Rodríguez-Espigares, I. et al. GPCRmd uncovers the dynamics of the 3D-GPCRome. Nat. Methods 17, 777–787 (2020).

14. Aranda-Garcia, D. et al. Simulating time-resolved dynamics of biomolecular systems. In Comprehensive Pharmacology 115–134 (Elsevier, 2022).

15. Suomivuori, C.-M. et al. Molecular mechanism of biased signaling in a prototypical G protein-coupled receptor. Science 367, 881–887 (2020).

16. Nivedha, A. K. et al. Identifying Functional Hotspot Residues for Biased Ligand Design in G-Protein-Coupled Receptors. Mol. Pharmacol. 93, 288–296 (2018).

17. Namkung, Y. et al. Monitoring G protein-coupled receptor and β-arrestin trafficking in live cells using enhanced bystander BRET. Nat. Commun. 7, 12178 (2016).

18. Capra, J. A. & Singh, M. Predicting functionally important residues from sequence conservation. Bioinformatics 23, 1875–1882 (2007).

19. Pándy-Szekeres, G. et al. GPCRdb in 2018: adding GPCR structure models and ligands. Nucleic Acids Res. 46, D440–D446 (2018).

20. Avet, C. et al. Effector membrane translocation biosensors reveal G protein and βarrestin coupling profiles of 100 therapeutically relevant GPCRs. Elife 11, e74101 (2022).

21. Isberg, V. et al. Generic GPCR residue numbers - aligning topology maps while minding the gaps. Trends Pharmacol. Sci. 36, 22–31 (2015).

22. Li, A., Liu, S., Huang, R., Ahn, S. & Lefkowitz, R. J. Loss of biased signaling at a G protein-coupled receptor in overexpressed systems. PLoS One 18, e0283477 (2023).

23. Heydenreich, F. M. et al. Molecular determinants of ligand efficacy and potency in GPCR signaling. Science 382, eadh1859 (2023).

24. Oakley, R. H., Laporte, S. A., Holt, J. A., Caron, M. G. & Barak, L. S. Differential affinities of visual arrestin, beta arrestin1, and beta arrestin2 for G protein-coupled receptors delineate two major classes of receptors. J. Biol. Chem. 275, 17201–17210 (2000).

25. Bhattacharya, S. & Vaidehi, N. Differences in allosteric communication pipelines in the inactive and active states of a GPCR. Biophys. J. 107, 422–434 (2014).

26. Ma, N., Nivedha, A. K. & Vaidehi, N. Allosteric communication regulates ligand-specific GPCR activity. FEBS J. 288, 2502–2512 (2021).

27. Laine, E., Auclair, C. & Tchertanov, L. Allosteric communication across the native and mutated KIT receptor tyrosine kinase. PLoS Comput. Biol. 8, e1002661 (2012).

28. Rivalta, I. et al. Allosteric Communication Disrupted by a Small Molecule Binding to the Imidazole Glycerol Phosphate Synthase Protein-Protein Interface. Biochemistry 55, 6484–6494 (2016).

29. Hegron, A. et al. Identification of Key Regions Mediating Human Melatonin Type 1 Receptor Functional Selectivity Revealed by Natural Variants. ACS Pharmacol Transl Sci 4, 1614–1627 (2021).

30. Al-Zoubi, R., Morales, P. & Reggio, P. H. Structural Insights into CB1 Receptor Biased Signaling. Int. J. Mol. Sci. 20, 1837 (2019).

31. Morales, P., Goya, P. & Jagerovic, N. Emerging strategies targeting CB cannabinoid receptor: Biased agonism and allosterism. Biochem. Pharmacol. 157, 8–17 (2018).

32. Wootten, D., Christopoulos, A., Marti-Solano, M., Babu, M. M. & Sexton, P. M. Mechanisms of signalling and biased agonism in G protein-coupled receptors. Nat. Rev. Mol. Cell Biol. 19, 638–653 (2018).

33. Cong, X. et al. Molecular insights into the biased signaling mechanism of the μ-opioid receptor. Mol. Cell 81, 4165–4175.e6 (2021).

34. Fenalti, G. et al. Molecular control of δ-opioid receptor signalling. Nature 506, 191–196 (2014).

35. Heydenreich, F. M. et al. High-throughput mutagenesis using a two-fragment PCR approach. Sci. Rep. 7, 6787 (2017).

36. Quoyer, J. et al. Pepducin targeting the C-X-C chemokine receptor type 4 acts as a biased agonist favoring activation of the inhibitory G protein. Proc. Natl. Acad. Sci. U. S. A. 110, E5088–97 (2013).

37. Beautrait, A. et al. Mapping the putative G protein-coupled receptor (GPCR) docking site on GPCR kinase 2: insights from intact cell phosphorylation and recruitment assays. J. Biol. Chem. 289, 25262–25275 (2014).

38. Beautrait, A. et al. A new inhibitor of the β-arrestin/AP2 endocytic complex reveals interplay between GPCR internalization and signalling. Nat. Commun. 8, 15054 (2017).

39. Li, X. et al. Crystal Structure of the Human Cannabinoid Receptor CB_2_. Cell 176, 459– 467.e13 (2019).

40. Hua, T. et al. Activation and Signaling Mechanism Revealed by Cannabinoid Receptor-G Complex Structures. Cell 180, 655–665.e18 (2020).

41. Lomize, M. A., Pogozheva, I. D., Joo, H., Mosberg, H. I. & Lomize, A. L. OPM database and PPM web server: resources for positioning of proteins in membranes. Nucleic Acids Res. 40, D370–6 (2012).

42. Jo, S., Kim, T., Iyer, V. G. & Im, W. CHARMM-GUI: a web-based graphical user interface for CHARMM. J. Comput. Chem. 29, 1859–1865 (2008).

43. Berendsen, H. J. C., Postma, J. P. M., van Gunsteren, W. F., DiNola, A. & Haak, J. R. Molecular dynamics with coupling to an external bath. J. Chem. Phys. 81, 3684–3690 (1984).

44. Harvey, M. J., Giupponi, G. & Fabritiis, G. D. ACEMD: Accelerating Biomolecular Dynamics in the Microsecond Time Scale. J. Chem. Theory Comput. 5, 1632–1639 (2009).

45. Huang, J. et al. CHARMM36m: an improved force field for folded and intrinsically disordered proteins. Nat. Methods 14, 71–73 (2017).

46. Vanommeslaeghe, K. & MacKerell, A. D., Jr. Automation of the CHARMM General Force Field (CGenFF) I: bond perception and atom typing. J. Chem. Inf. Model. 52, 3144–3154 (2012).

47. Vanommeslaeghe, K., Raman, E. P. & MacKerell, A. D., Jr. Automation of the CHARMM General Force Field (CGenFF) II: assignment of bonded parameters and partial atomic charges. J. Chem. Inf. Model. 52, 3155–3168 (2012).

48. Vanommeslaeghe, K. et al. CHARMM general force field: A force field for drug-like molecules compatible with the CHARMM all-atom additive biological force fields. J. Comput. Chem. 31, 671–690 (2010).

49. Yu, W., He, X., Vanommeslaeghe, K. & MacKerell, A. D., Jr. Extension of the CHARMM General Force Field to sulfonyl-containing compounds and its utility in biomolecular simulations. J. Comput. Chem. 33, 2451–2468 (2012).

50. Foutch, D., Pham, B. & Shen, T. Protein conformational switch discerned via network centrality properties. Comput. Struct. Biotechnol. J. 19, 3599–3608 (2021).

