## Supplementary Information for "Multiple triggers converge to preferential effector coupling in the CB_2_R through a complex allosteric communication network"

### Table of content

Supplementary note
Supplementary tables
Supplementary figures

### Supplementary note

**Note 1 - Detailed structural description of PrefCoup$_{G\alpha i2}$ cluster 1 to 3**

In the PrefCoup$_{G\alpha i2}$ cluster 1, logistic regression highlights differences in contact stability and dynamics for three conserved domains (**Figure 5B**): (1) The conserved W258$^{6x48}$, also known as the toggle switch, within the CWxP motif in TM6 shows an increased interaction frequency with residue L195$^{5x44}$ in Gαi2 preferential mutants. (2) Two residues in the sodium binding site (D80$^{2x50}$ and S120$^{3x39}$) show an increased interaction stability in Gαi2 preferential mutants. (3) Finally, we also find differences involving the E/DRY motif in the intracellular half of TM3. Two consecutive residues from this motif, R131$^{3x50}$ and Y132$^{3x51}$, show a decreased interaction stability with residue V212$^{5x61}$ in Gαi2 preferential mutants.

Interestingly, in the PrefCoup$_{G\alpha i2}$ cluster 2 (**Figure 5C**), we identify alterations in contact dynamics for the same motifs as in cluster 1, but with different residues implicated: (1) The toggle switch (W258$^{648}$) is involved in two interactions with C288$^{7x41}$ and N291$^{7x45}$ that lose stability in Gαi2 preferential mutants. The latter residues are also part of the sodium binding site. (2) In the sodium binding site (besides contacts with N291$^{7x45}$), residue S292$^{7x46}$ is establishing contacts with S47$^{1x46}$ which are more stable in Gαi2 preferential mutants. (3) Finally, the DRY motif shows a modified interaction involving residues R131$^{3x50}$ and T127$^{3x46}$. This interaction is established between stacked residues in the helical structure of TM3 and is more stable in Gαi2 preferential mutants. Moreover, we have identified a densely populated region of highlighted interactions in the extracellular halves of TM7, 1, and 2, comprising residues C40$^{1x39}$, F91$^{2x61}$, F87$^{2x57}$, S285$^{7x38}$, and G44$^{1x43}$. Generally, the interactions within this network are less stable in Gαi2 preferential mutants, with only one interaction showing increased stability, i.e. between residue C40$^{1x39}$ and F87$^{2x57}$.

In PrefCoup$_{G\alpha i2}$ cluster 3 (**Figure 5D**), the logistic regression model highlights differences in the NPxxY motif (1) and the sodium binding site (2). The NPxxY motif stands out with two highlighted inter-residual contacts that are more stable in Gαi2 preferential mutants. One interaction is with residue N51$^{1x50}$, and the other is with residue D80$^{2x50}$ in the sodium binding site. (3) The DRY motif also shows altered interactions in this cluster. Specifically, we observe two interactions involving residue R131$^{3x50}$ that are more stable in Gαi2 preferential mutants and are established with residues T127$^{3x46}$ and T246$^{6x36}$.

### Supplementary tables

Table S1. Simulated mutants indicating the sequence position, GPCRdb notation, coupling profile classification

| Position | GPCRdb notation | Coupling profile |
|---|---|---|
| 33 | 1x32 | Coup_Gi_bArr |
| 47 | 1x46 | Coup_Gi_bArr |
| 49 | 1x48 | Coup_Gi_bArr |
| 52 | 1x51 | Coup_Gi_bArr |
| 53 | 1x52 | Coup_Gi_bArr |
| 61 | 1x60 | PrefCoup_Gi |
| 77 | 2x47 | PrefCoup_Gi |
| 91 | 2x61 | Coup_Gi_bArr |
| 109 | 3x28 | PrefCoup_Gi |
| 112 | 3x31 | Coup_Gi_bArr |
| 117 | 3x36 | PrefCoup_Gi |
| 119 | 3x38 | Coup_Gi_bArr |
| 121 | 3x40 | Coup_Gi_bArr |
| 125 | 3x44 | PrefCoup_Gi |
| 152 | 4x44 | Coup_Gi_bArr |
| 159 | 4x51 | Coup_Gi_bArr |
| 165 | 4x57 | Coup_Gi_bArr |
| 176 | 176 | PrefCoup_Gi |
| 178 | 178 | Coup_Gi_bArr |
| 185 | 185 | Coup_Gi_bArr |
| 199 | 5x48 | PrefCoup_Gi |
| 203 | 5x52 | Coup_Gi_bArr |
| 205 | 5x54 | PrefCoup_Gi |
| 217 | 5x66 | PrefCoup_Gi |
| 251 | 6x41 | Coup_Gi_bArr |
| 282 | 7x35 | Coup_Gi_bArr |
| 285 | 7x38 | PrefCoup_Gi |
| 291 | 7x45 | PrefCoup_Gi |
| 292 | 7x46 | PrefCoup_Gi |
| 293 | 7x47 | Coup_Gi_bArr |
| 297 | 7x51 | PrefCoup_Gi |
| 302 | 7x56 | PrefCoup_Gi |
| 313 | 8x57 | Coup_Gi_bArr |
| 319 | 8x63 | Coup_Gi_bArr |

### Supplementary figures

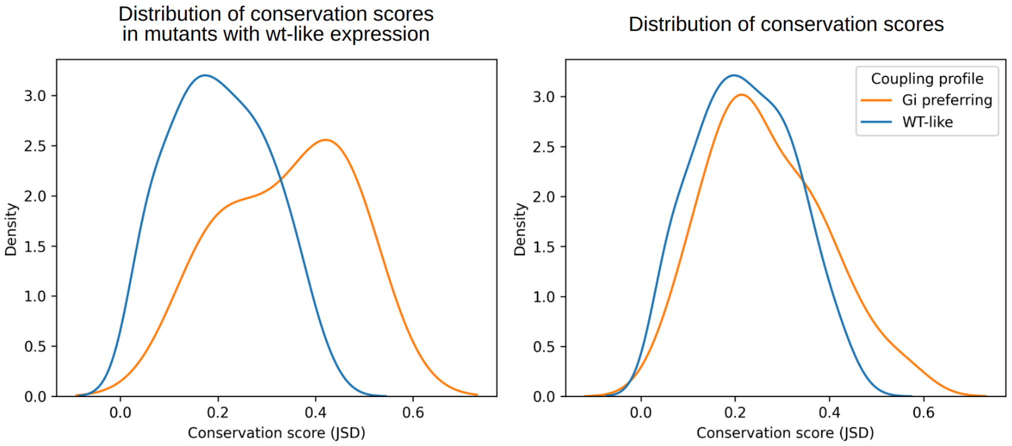


**Figure S1. Evolutionary conservation in mutants that don't affect expression.** Comparison of the distribution of conservation on coupling profiles of interest between residues that do not impact expression against all the residues. Conservation scores are computed using the Jensen Shannon divergence on an MSA of class A GPCRs.

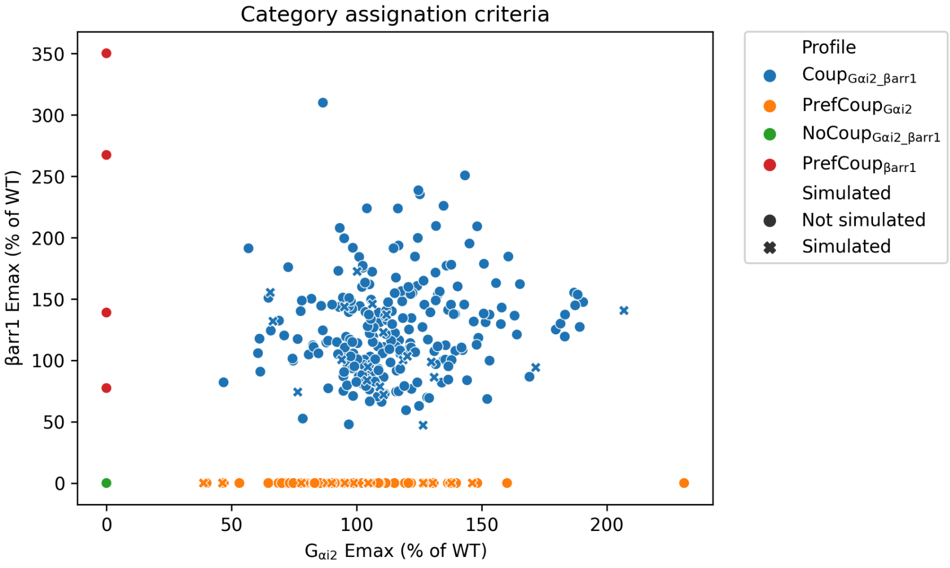


**Figure S2. Coupling profile categorization criteria.** Scatterplot showing the Emax distribution of mutants for Gαi2 and βarr1 coupling. Mutants are coloured based on their coupling profile as described in Figure. A large fraction of mutants with preserved coupling ($Coup_{G\alpha i\_\beta arr1}$) and Gi preferential mutants ($PrefCoup_{G\alpha i}$) have been simulated. Those mutants are represented with a dot. Mutants highlighted with an 'x' are not simulated as they do not fulfill the following conditions: (1) cell surface expression similar to the WT level and (2) located in receptor region with a solved 3D structure.

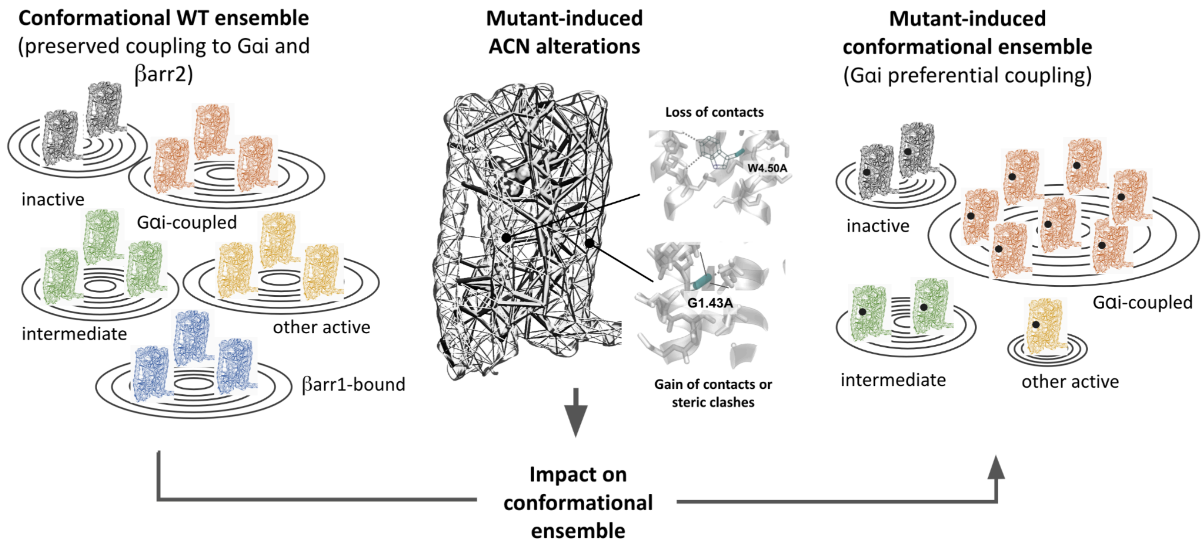


**Figure S3**. Schematic model of mutant-induced alterations of allosteric communication networks (ACN) and their impact on the conformational ensemble comprising inactive, intermediate, and various active states.

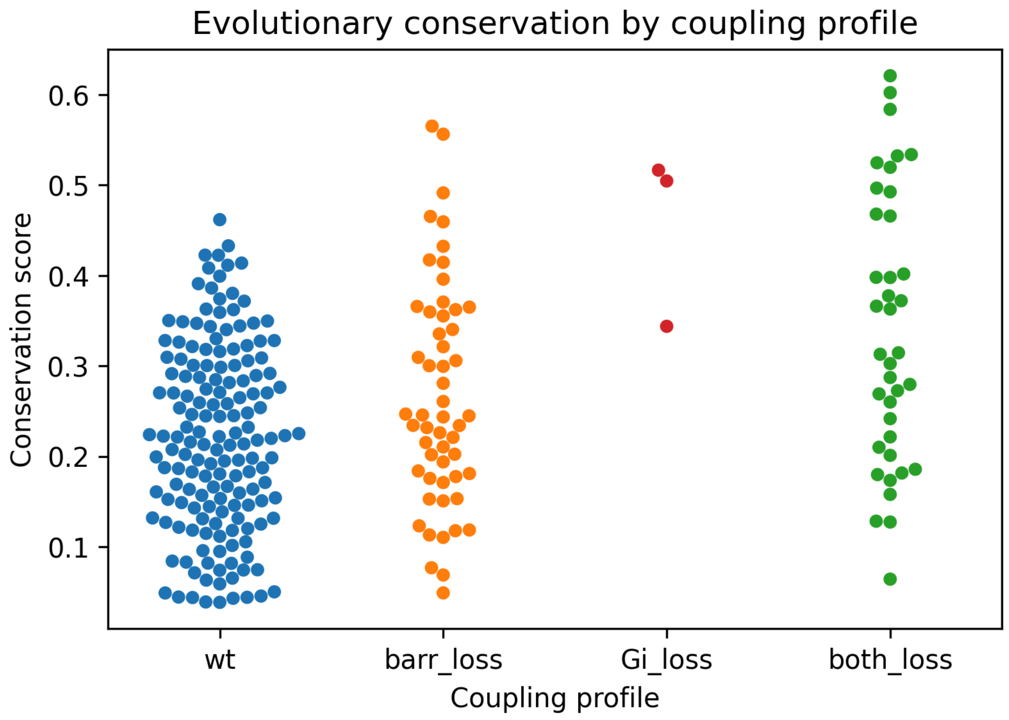


**Figure S4. Evolutionary conservation by coupling profile.** This swarm plot shows the distribution of conservation scores in each one of the coupling profiles. The conservation score is computed as the Jensen Shannon Divergence of a class A GPCR multiple sequence alignment.

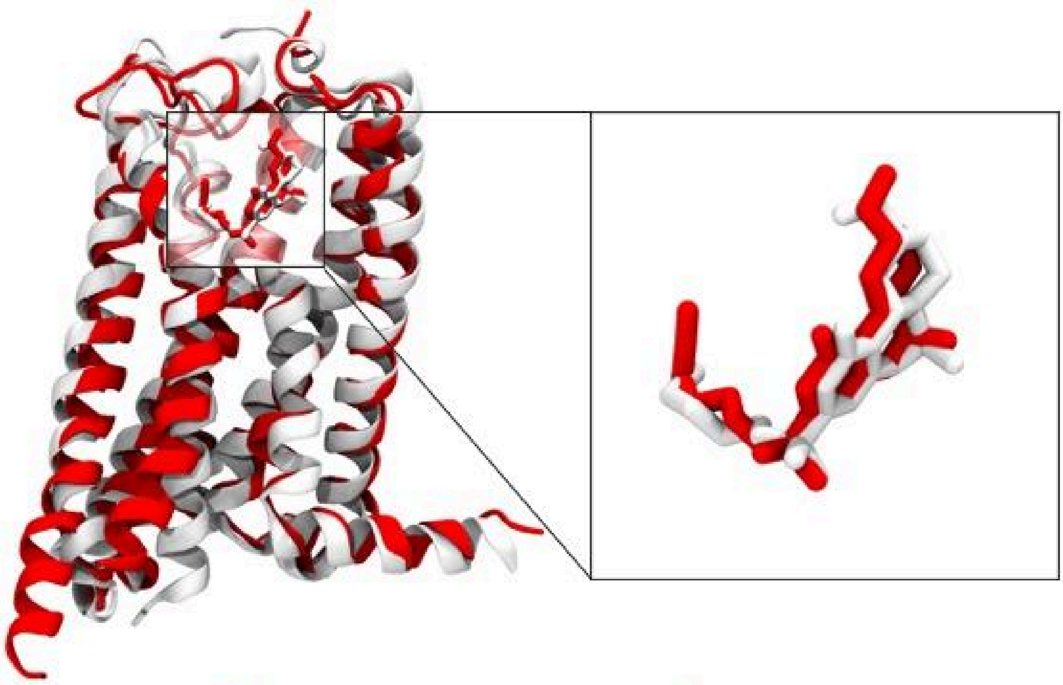

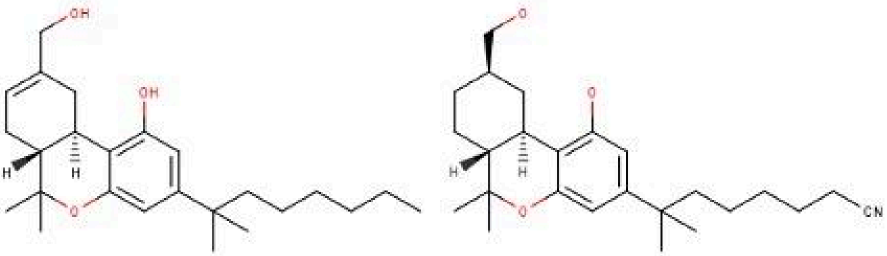


HU-210 AM12033

**Figure S5. Comparison of CB2 Receptor (CB2R) Conformations and Ligand Docking Models.** The white ribbon diagram represents the inactive CB2R structure modeled from PDB entry 5ZTY, while the red ribbon diagram overlaid represents the active CB2R conformation from PDB entry 6KPC. Highlighted within the receptor is the compound HU-210 docked into the inactive structure, positioned to mimic the binding pose of the red molecule, AM12033, which is docked in the active structure. Below are the 2D chemical structures of HU-210 and AM12033, illustrating their molecular resemblance.

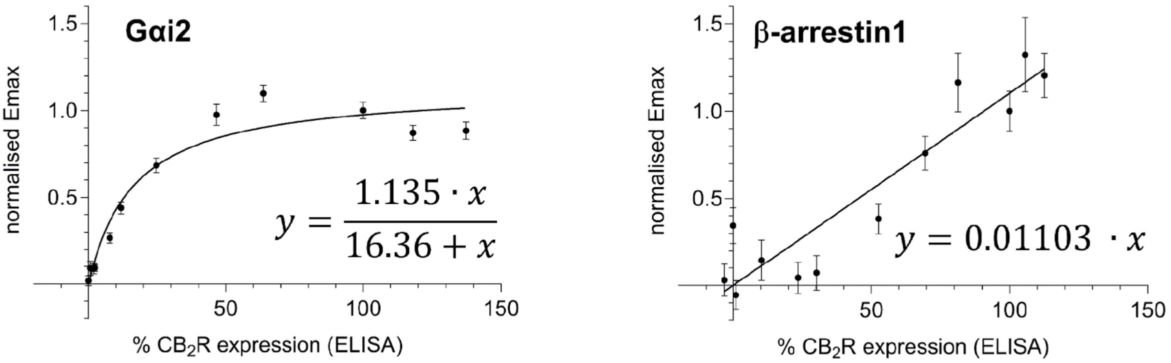


**Figure S6**. Correlation of $CB_2R$ expression levels with measured Emax values, used to calculate the correction factors in Supplementary Data 1.
